# Organism-level visual genotyping of knockout zygosity enables functional gene analysis and modular combination with reporter transgenes

**DOI:** 10.64898/2026.01.16.699911

**Authors:** Franziska Krämer, Julia Ratke, Frederic Strobl

**Author notes:** these authors contributed equally (Franziska Krämer, Julia Ratke). **Corresponding Author:** Frederic Strobl.

## Abstract

Analysis of gene function in diploid organisms typically relies on molecular methods to distinguish wild-type, mono-allelic knockout, and bi-allelic knockout individuals, creating a major practical bottleneck for large-scale developmental studies. Here, we establish a modular visual genotyping approach that enables organism-level discrimination of knockout zygosity by tagging alternative disrupted alleles with spectrally distinct fluorescent markers. Implemented in the red flour beetle *Tribolium castaneum*, a two-marker strategy permits reliable identification of bi-allelic knockouts within mixed cohorts in a semi-random insertional mutagenesis framework. Further, a four-marker strategy using a targeted CRISPR/Cas9-based gene editing approach enables systematic generation and unambiguous recognition of individuals homozygous for both a disrupted gene of choice and a fluorescent reporter transgene. This design substantially reduces the workload associated with routine molecular genotyping while enabling genotype-resolved long-term live imaging of embryonic morphogenesis. Application to the extra-embryonic specification factor *zerknüllt 1* reveals haplosufficiency and a semi-lethal phenotype associated with variable morphogenetic outcomes. Together, our approach provides a scalable framework for functional analysis of essential developmental genes.

**Summary Statement:** Visual marker-based genotyping enables organism-level discrimination of knockout zygosity, reducing reliance on molecular assays and facilitating scalable, genotype-resolved live imaging of developmental gene function in diploid model organisms.

## Introduction

Analysis of gene function fundamentally relies on phenotypic characterization of loss-of-function conditions. In multicellular diploid organisms, including widely used developmental models such as mice (Capecchi, 2005), zebrafish (Haffter et al., 1996) and insects (St Johnston, 2002), random or targeted gene disruption typically produces mono-allelic knockout individuals (hetero- or hemizygotes) that retain one functional allele and therefore often show no or only partial phenotypes. Functional interpretation therefore requires analysis of bi-allelic knockout individuals (homozygotes) lacking all gene activity. These are commonly generated by mating mono-allelic carriers, yielding mixed progeny composed of wild-type (25%), mono-allelic knockout (50%), and bi-allelic knockout (25%) individuals (Coleman et al., 2015).

Genotype inference based solely on morphology is often unreliable. Phenotypes may be subtle, variably penetrant (Raj et al., 2010), influenced by genetic background (Nadeau, 2001) or environmental conditions (Gilbert, 2017), or induced by experimental manipulation. Consequently, phenotype-based genotype assignment is prone to both false-negative and false-positive errors that can propagate through datasets and bias genotype-phenotype relationships. These risks can only be mitigated through independent genotype determination.

This requirement represents a major practical bottleneck in developmental biology. Quantitative phenotypic analyses depend on large cohorts, biological replicates, and statistically powered comparisons, making genotype determination a substantial manpower constraint. The need to genotype each individual using molecular methods – including PCR-based assays or sequencing – leads to rapidly accumulating demands on labor, cost, and time. These constraints are further amplified in multi-generational crosses intended to combine two or more genetic modifications (St Johnston, 2002), as Mendelian segregation reduces recovery rate of desired genotypes with each additional locus. Even when gene disruptions include visible transformation markers – such as coat color-based markers in mouse (Zheng et al., 1999), fluorescent heart-or lens-specific markers in zebrafish (Kwan et al., 2007), or the widely used 3×P3-driven fluorescent eye markers in insects (Berghammer et al., 1999) – genotype discrimination is generally limited to transgene presence versus absence. Consequently, zygosity often remains unresolved, necessitating molecular genotyping.

To partially address this limitation, we previously established the AGameOfClones (AGOC) and AClashOfStrings (ACOS) vector concepts, which employ spectrally distinct fluorescent eye markers based on mOrange and mCherry (mO and mC) respectively mCerulean and mVenus (mCe and mVe) to enable organism-level genotyping of up to two transgenes. The functionality of both concepts was demonstrated in the red flour beetle *Tribolium castaneum*, an emerging model for mechanistic studies of embryogenesis (Brown et al., 2009), through the systematic generation of single and double homozygous reporter lines (Strobl et al., 2018; Strobl et al., 2023).

We reasoned that this strategy could be extended to gene knockout analysis. By generating two alternative disrupted alleles of a gene, each tagged with a visually distinguishable marker, wild types, mono-allelic knockouts and bi-allelic knockouts could in principle be discriminated directly at the organismal level. Such a design preserves the gold-standard framework of knockout analysis – comparative assessment of all genotypes within a single cohort – while reducing reliance on molecular genotyping.

Here, we establish and validate this approach in *Tribolium*. We first implement a two-marker strategy in a semi-random insertional mutagenesis framework and demonstrate unambiguous organism-level discrimination among genotypes. We then extend the concept to a four-marker configuration that enables simultaneous determination of knockout and reporter zygosity, thereby facilitating genotype-resolved functional live imaging workflows. Finally, we discuss the advantages, limitations, and potential future refinements of our approach.

## Results

### A two-marker strategy for gene function analysis

To test whether knockout genotypes can be identified visually through complementary disrupted alleles distinguished by spectrally distinct fluorescence, we implemented a two-marker strategy in *Tribolium* within a piggyBac-based semi-random insertional mutagenesis framework inspired by a previous large-scale screen (Trauner et al., 2009).

At first, we designed an AGOC/ACOS-inspired construct containing mVe and mO within interwoven LoxP/LoxN site pairs that allow mutually exclusive Cre-mediated marker excision (Figure 1A). Embryonic injections yielded multiple F1 individuals potentially mosaic for germline transformation. From five independent outcrosses, we recovered F2 progeny carrying both transformation markers. For each transgenic lineage, a single F2 founder was selected for further outcrossing. Segregation analysis confirmed Mendelian inheritance consistent with single-locus insertions (Supplementary Table 1, ‘F2’ row), allowing immediate establishment of five independent sublines (#1–#5). To generate complementarily marked alleles for subsequent combination into visually identifiable genotypes, we implemented a defined mating procedure involving the pFIRE{HSP68’NLS-Cre} #1 helper subline, which expresses a nuclear-localized Cre recombinase and carries mC as a transformation marker (Strobl and Stelzer, 2021). The procedure comprised three mating steps across four generations:

1. F3 (mVe-mO) hemizygotes were mated with (mC) hemizygotes of the helper subline (Figure 1B, ‘F3’ row), which resulted in F4 (mC; mVe-mO) double hemizygotes (Supplementary Table 1, ‘F3’ row) in which Cre-mediated recombination occurs.
2. F4 (mC; mVe-mO) double hemizygotes were mated with wild types (Figure 2, ‘F4’ row). Due to recombination in the germline, this resulted in F5 (mVe) as well as (mO) post-recombination hemizygotes (Supplementary Table 1, ‘F4’ row).
3. F5 (mVe) hemizygotes were mated with F5 (mO) hemizygotes (Figure 2, ‘F5’ row), which resulted in ∼25% F6 (mVe/mO) heterozygotes (Supplementary Table 1, ‘F5’ row). In cases where the insertion disrupted a gene, these individuals inherited two disrupted alleles and thus represented bi-allelic knockouts.

**Figure 1.**
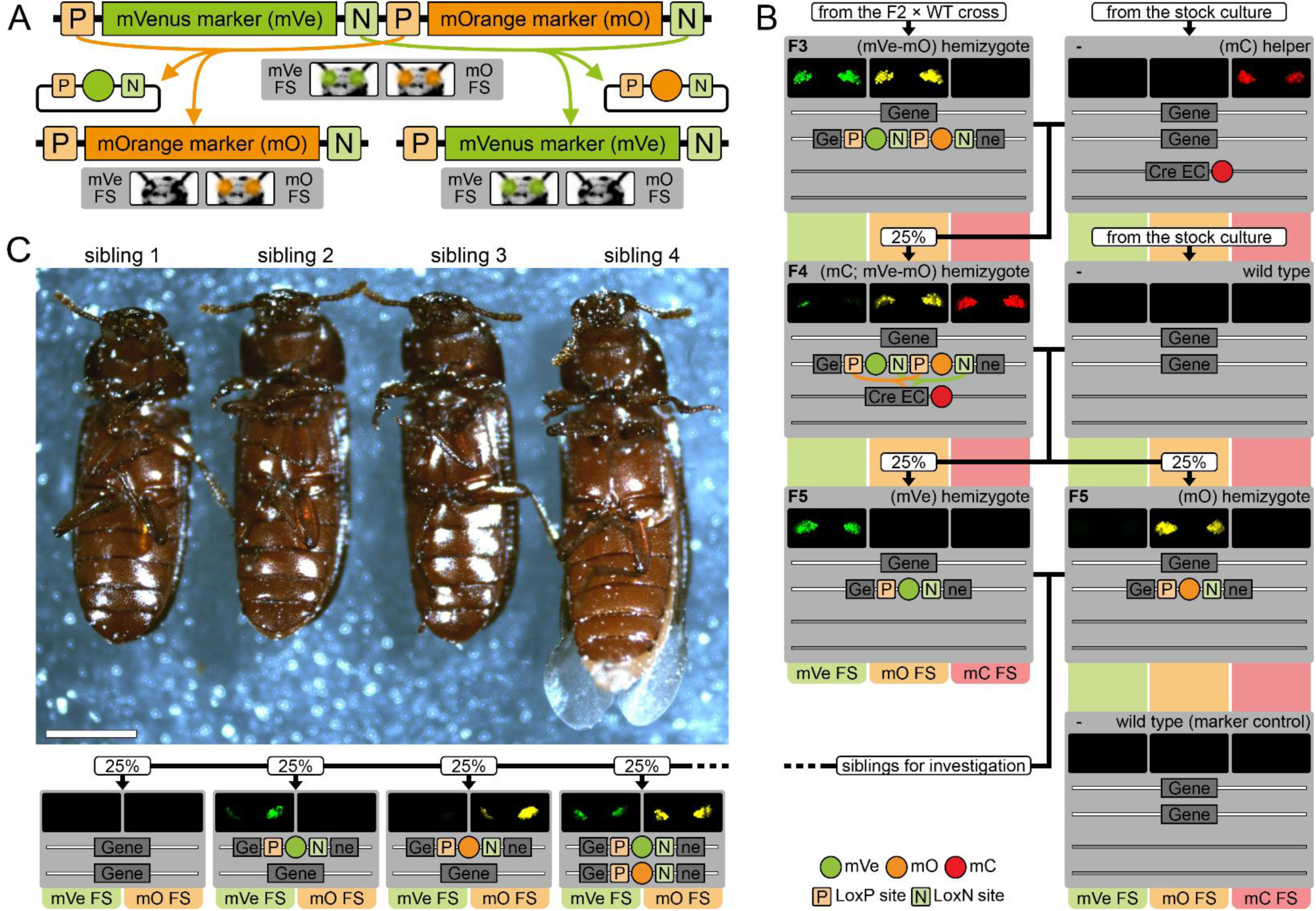
| A two-marker strategy for gene function analysis. **(a)** Genetic construct used for semi-random insertional mutagenesis. Two fluorescent eye markers are embedded within interwoven Lox site pairs, allowing mutually exclusive Cre-mediated excision of one marker. Gray boxes indicate marker phenotypes before and after recombination. **(b)** Mating procedure for a systematic generation of bi-allelic knockout individuals in cases where the insertion disrupted a gene. Phenotypic documentation for the #1 subline. **(c)** Four male siblings from the #5 subline derived from a cross between (mVe) and (mO) hemizygotes. Gray boxes show visual markers (pseudo-colored) and inferred genotypes. All specimens were alive at the time of imaging. P, LoxP site; N, LoxN site; FS, filter set; WT, wild type; Cre, expression cassette for the Cre recombinase.

**Figure 2.**
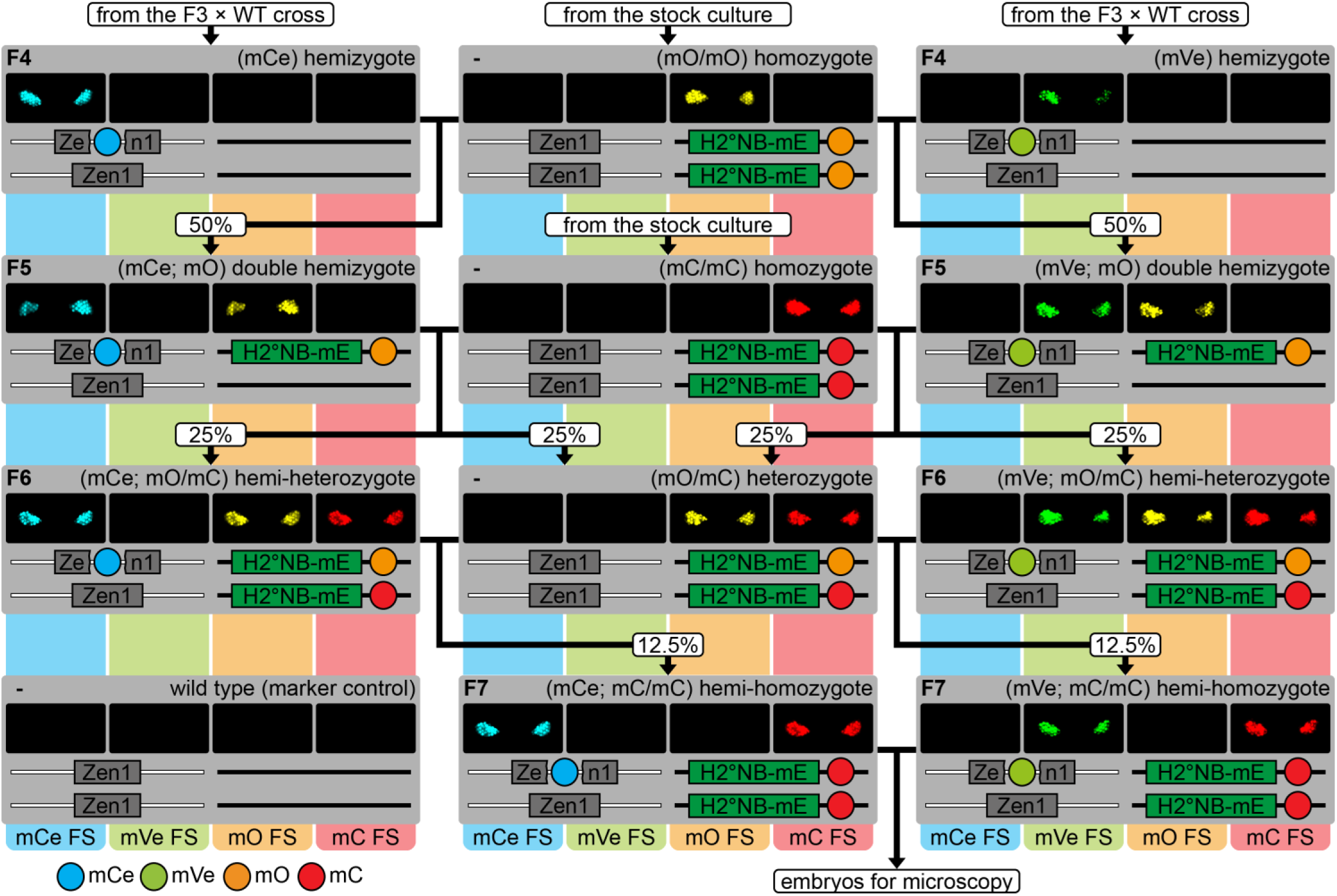
| Mating procedure for a systematic generation of knockout-reporter genotypes. Phenotypic documentation for replicate 1. Gray boxes show visual markers (pseudo-colored) and inferred genotypes. Rounded rectangles indicate the theoretical Mendelian progeny ratios. WT, wild type; H2°NB-mE, expression cassette for mEmerald-labeled anti-histone nanobodies; FS, filter set.

In four sublines (#1–#4), F6 siblings of all genotypes were morphologically indistinguishable. In contrast, subline #5 displayed a clear genotype-phenotype association: siblings lacking one or both markers showed normal morphology, whereas inheritance of both mVe and mO consistently co-segregated with a pronounced ‘terminal defect’ phenotype. (Figure 1C; Supplementary Figures 1 and 2). This pattern is consistent with haplosufficiency of the disrupted gene, with phenotypic manifestation only in the absence of any functional allele. Importantly, strict phenotype-marker co-segregation demonstrates that the two-marker strategy enables reliable organism-level identification of bi-allelic knockouts based solely on marker scoring.

### A four-marker strategy for gene function analysis using fluorescence microscopy

Following validation of the two-marker approach, we extended the framework to a four-marker strategy by incorporating fluorescent reporter transgenes to enable morphogenetic analysis of knockout phenotypes by fluorescence microscopy, analogous to our previous combination of two reporter constructs (Strobl and Stelzer, 2021; Strobl et al., 2023).

To demonstrate applicability for functional analysis during embryogenesis, we transitioned from semi-random to site-specific insertional mutagenesis and targeted *zerknüllt 1* (*Zen1*, NCBI Gene ID: 641533), a transcription factor required for extra-embryonic membrane specification (Jain et al., 2020; Mann and Panfilio, 2024; Panfilio et al., 2013; van der Zee et al., 2005).

At first, we designed a CRISPR/Cas9-based gene editing approach to independently insert either mCe or mVe into the *Zen1* coding sequence via homology-directed repair, thereby introducing a premature stop codon at the beginning of the third exon (Supplementary Figure 3). Embryonic injections yielded multiple F1 individuals potentially mosaic for germline transformation. From subsequent outcrosses, we recovered one F2 founder carrying mCe and one F2 founder carrying mVe, both of which were further outcrossed. Among the F3 progeny, three transgenic descendants per founder were selected for further outcrossing. Segregation analysis demonstrated Mendelian inheritance consistent with a single-locus insertion for each of the six individuals (Supplementary Tables 2 and 3, ‘F3’ row).

To generate knockout-reporter hybrids with suitable genotypes, we implemented a defined mating procedure involving the transgenic AGOC{ATub’H2A/H2B°NB-mEmerald} #1 reporter subline, which expresses mEmerald-labeled anti-histone nanobodies and is available in both the (mO/mO) and (mC/mC) ‘flavor’ (Strobl et al., 2025). The procedure was performed in three biological replicates, each using one F4 (mCe) and one F4 (mVe) individual, and comprised four mating steps across five generations:

1. F4 (mCe) as well as (mVe) hemizygous knockouts were separately mated with (mO/mO) homozygotes of the reporter subline (Figure 2, ‘F4’ row). This resulted in F5 (mCe; mO) and (mVe; mO) double hemizygotes (Supplementary Tables 2 and 3, ‘F4’ rows).
2. F5 (mCe; mO) as well as (mVe; mO) double hemizygotes were separately mated with (mC/mC) homozygotes of the same reporter subline (Figure 2, ‘F5’ row), which resulted in F6 hemi-heterozygotes carrying three markers each (Supplementary Tables 2 and 3, ‘F5’ rows).
3. F6 (mCe; mO/mC) as well as (mVe; mO/mC) hemi-heterozygotes were separately mated with (mO/mC) heterozygous siblings (Figure 2, ‘F6’ row), which resulted in F7 (mCe; mC/mC) as well as (mVe; mC/mC) hemi-homozygotes (Figure 2 and Supplementary Tables 2 and 3, ‘F6’ rows). To confirm these two genotypes, respective individuals were outcrossed (Supplementary Tables 2 and 3, ‘F7C’ rows).
4. F7 (mCe; mC/mC) hemi-homozygotes were mated with (mVe; mC/mC) hemi-homozygotes (Figure 2, ‘F7’ row), which resulted in (i) (mC/mC) homozygotes, (ii) (mCe; mC/mC) hemi-homozygotes, (iii) (mVe; mC/mC) hemi-homozygotes, and (iv) (mCe/mVe; mC/mC) hetero-homozygotes (which inherited two disrupted alleles and thus represent bi-allelic *Zen1* knockouts).

From the F3 to the F6 generation, marker distributions were consistent with the theoretical Mendelian ratios (Supplementary Tables 2 and 3, ‘*p*’ column). In contrast, in the F7 generation, progeny ratios differed significantly from expectations in all three replicates (Table 1, ‘F7’ row), with a pronounced underrepresentation of (mCe/mVe; mC/mC) hetero-homozygotes. Despite this deviation, the descendants showed no obvious morphological differences from their siblings during larval, pupal and adult stages. To confirm genotype assignment, three adult females and thee adult males were selected for further outcrossing. Segregation analysis revealed Mendelian inheritance consistent with the predicted genotype (Table 1, ‘F8’ row), confirming that (mCe/mVe; mC/mC) individuals indeed represent bi-allelic knockouts for *Zen1*.

**Table 1.**
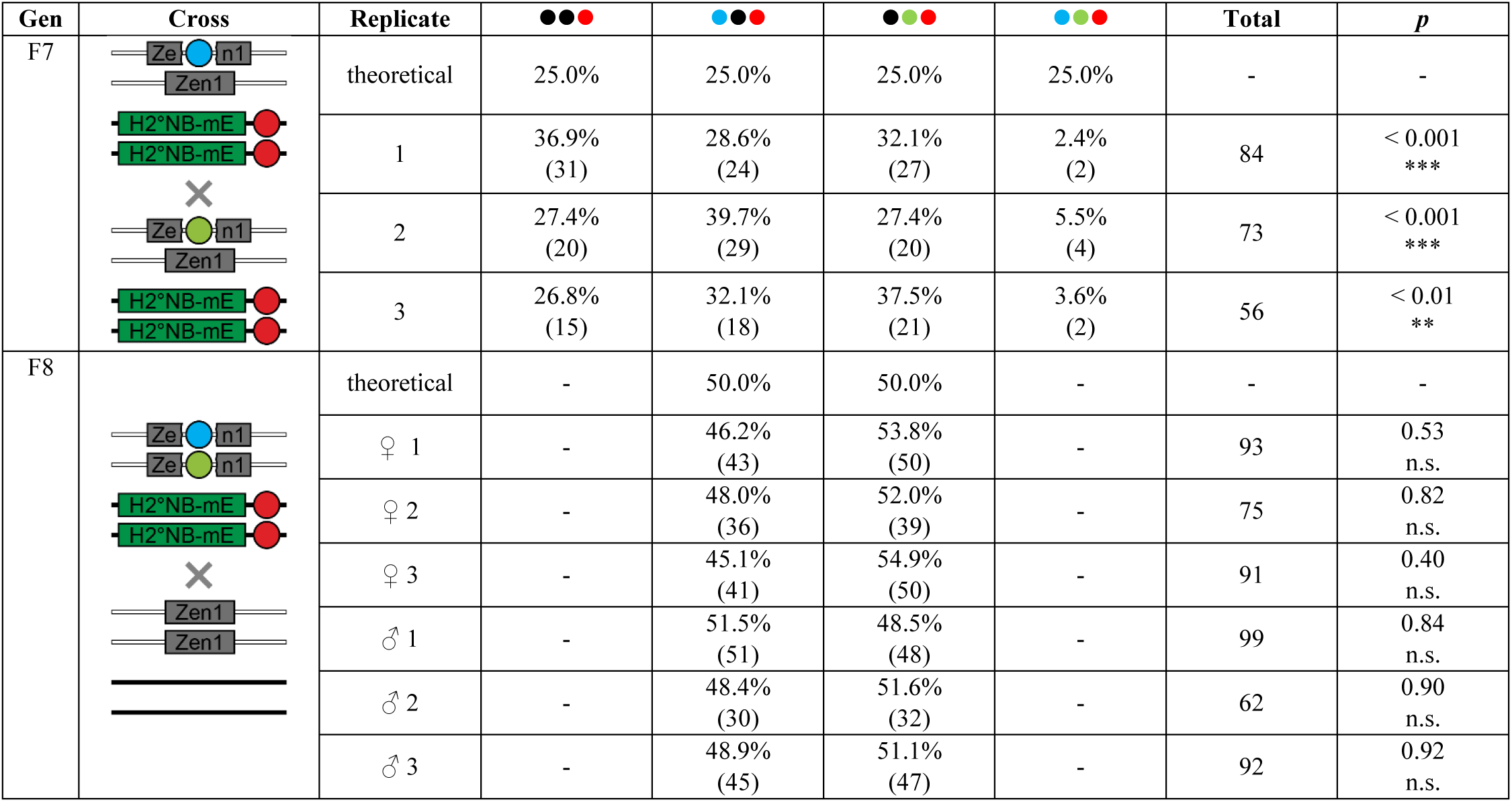
| F7 and F8 crossing results across three biological replicates. Progeny genotype frequencies were scored during the pupal stage and compared to the theoretical Mendelian ratio using a chi-square goodness-of-fit test. Percentages and numbers indicate observed genotype frequencies and absolute progeny scores, respectively. In the F7 cross, significant deviations from the theoretical ratios were found for all three repetitions, yet all four expected marker phenotypes were found. In the F8 control cross, no significant deviations were found. H2°NB-mE, expression cassette for mEmerald-labeled anti-histone nanobodies; n.s., not significant.

| Gen | Cross | Replicate | ●●● | ●●● | ●●● | ●●● | Total | *p* |
| --- | --- | --- | --- | --- | --- | --- | --- | --- |
| F7 |  | theoretical | 25.0% | 25.0% | 25.0% | 25.0% | - | - |
|  |  | 1 | 36.9%<br>(31) | 28.6%<br>(24) | 32.1%<br>(27) | 2.4%<br>(2) | 84 | < 0.001<br>*** |
|  |  | 2 | 27.4%<br>(20) | 39.7%<br>(29) | 27.4%<br>(20) | 5.5%<br>(4) | 73 | < 0.001<br>*** |
|  |  | 3 | 26.8%<br>(15) | 32.1%<br>(18) | 37.5%<br>(21) | 3.6%<br>(2) | 56 | < 0.01<br>** |
| F8 |  | theoretical | - | 50.0% | 50.0% | - | - | - |
|  |  | ♀ 1 | - | 46.2%<br>(43) | 53.8%<br>(50) | - | 93 | 0.53<br>n.s. |
|  |  | ♀ 2 | - | 48.0%<br>(36) | 52.0%<br>(39) | - | 75 | 0.82<br>n.s. |
|  |  | ♀ 3 | - | 45.1%<br>(41) | 54.9%<br>(50) | - | 91 | 0.40<br>n.s. |
|  |  | ♂ 1 | - | 51.5%<br>(51) | 48.5%<br>(48) | - | 99 | 0.84<br>n.s. |
|  |  | ♂ 2 | - | 48.4%<br>(30) | 51.6%<br>(32) | - | 62 | 0.90<br>n.s. |
|  |  | ♂ 3 | - | 48.9%<br>(45) | 51.1%<br>(47) | - | 92 | 0.92<br>n.s. |

Methodologically, these findings demonstrate that the four-marker strategy enables reliable recovery of hemi-homozygous knockout-reporter individuals, which in turn allows efficient generation of progeny bi-allelic for the disrupted gene and homozygous for the fluorescence expression cassette. Biologically, the altered genotype ratios among the F7 progeny indicate that biallelic disruption of *Zen1* reduces viability without causing complete lethality, indicative of a ‘semi-lethal’ effect (Trauner et al., 2009).

### Phenotypic analysis of the *Zen1* knockout based on long-term fluorescence live imaging

To investigate the biological basis of the observed semi-lethality, we focused on embryonic morphogenesis. This decision was guided by two observations: first, *Zen1* functional knockouts were morphologically inconspicuous during larval, pupal, and adult stages; second, previous studies have shown that *Zen1* expression is largely restricted to early embryogenesis.

We therefore set up mass crosses of (mCe; mC/mC) hemi-homozygous females and (mVe; mC/mC) hemi-homozygous males and subjected five embryos to long-term live imaging using light sheet fluorescence microscopy. Recordings were performed using acquisition parameters previously shown to permit non-invasive imaging of homozygous embryos from the AGOC{ATub’H2A/H2B°NB-mEmerald} #1 reporter subline alone (Strobl et al., 2025). Since this reporter line forms part of the genetic background analyzed here, using identical conditions minimized the likelihood that any aberrant phenotype arose from imaging-related perturbation rather than from disruption of *Zen1*.

Imaging commenced at the uniform blastoderm stage. All five embryos showed comparable mEmerald fluorescence levels throughout the recording period, consistent with homozygosity for the nanobody-expressing reporter transgene. However, imaging revealed two distinct classes of morphogenetic phenotypes. Three embryos developed normally (Figure 3, embryos 1–3; Supplementary Movies 1–3), resembling previously recorded development of the reporter subline alone. These embryos were retrieved after imaging, hatched successfully, and matured into fertile adults.

**Figure 3.**
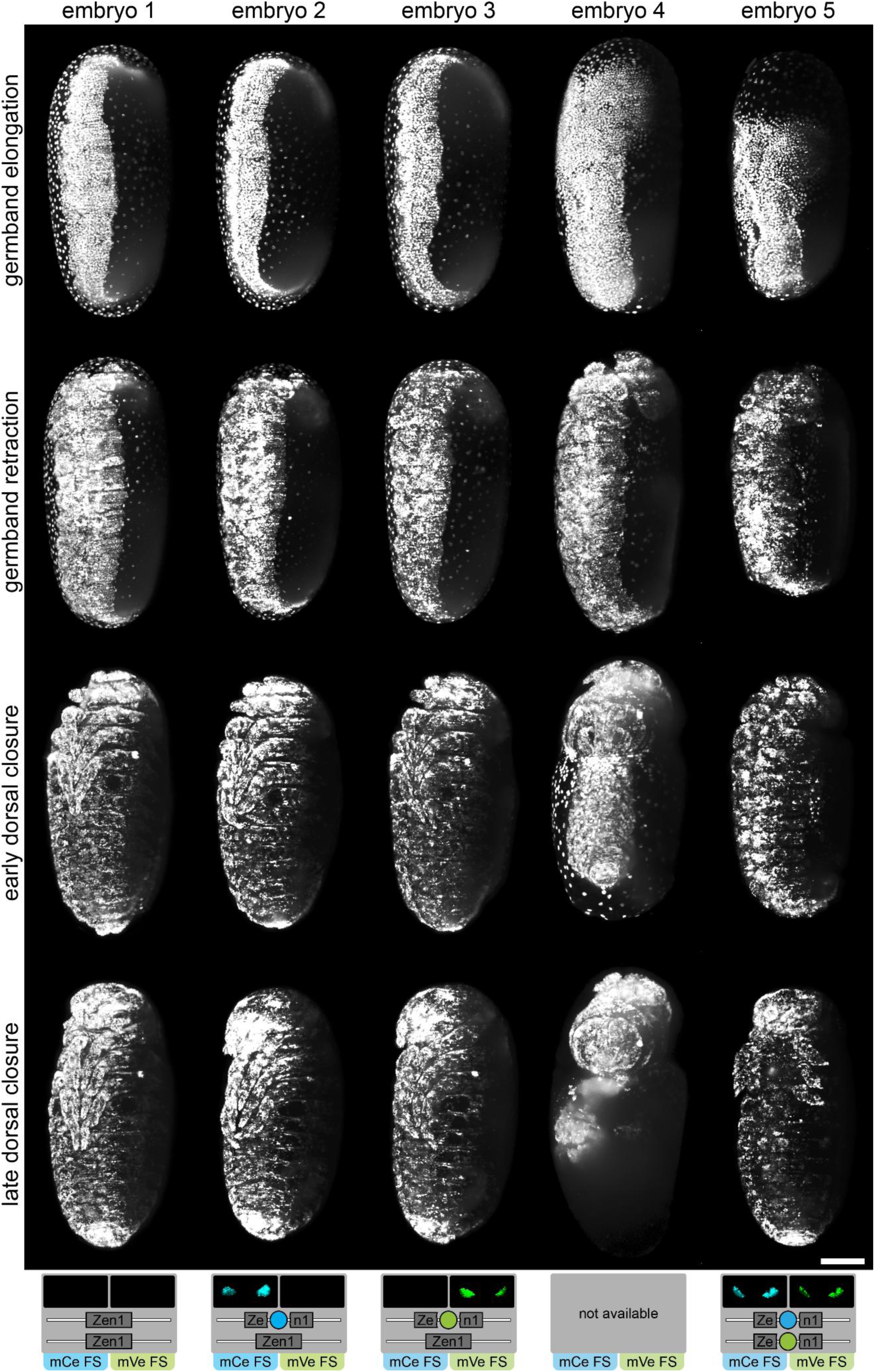
| Fluorescent reporter-assisted analysis of *Zen1* knockout phenotypes. Long-term live imaging of five embryos from the *Zen1* knockout-reporter background using light sheet fluorescence microscopy. Specimens are shown in the ventrolateral view as maximum projections. Gray boxes show visual markers (assessed after retrieved embryo were reared to adulthood, pseudo-coloring) and inferred genotypes. FS, filter set.

In contrast, two embryos displayed aberrant morphogenesis closely resembling the previously described *Zen1* knockdown phenotypes (Jain et al., 2020; Mann and Panfilio, 2024; Panfilio et al., 2013; van der Zee et al., 2005). In both cases, serosa window closure remained incomplete, leaving the germband in persistent contact with the vitelline envelope throughout germband elongation and retraction (Figure 3, ‘germband elongation’ and ‘germband retraction’ rows). During early dorsal closure, two distinct developmental outcomes were observed. One embryo showed an ‘insufficient withdrawal’ phenotype, in which reduced serosal movement caused the actomyosin cable – normally involved in dorsal organ formation – to constrict the embryo (Figure 3, embryo 4; Supplementary Movie 4). Although retrieved after imaging, this embryo failed to hatch. The other embryo displayed a ‘sufficient withdrawal’ phenotype, in which the serosa was able to pass over the posterior pole and completed a provisional mode of dorsal closure lacking clear dorsal organ formation. Additional abnormalities were observed, including outwardly oriented legs (Figure 3, embryo 5; Supplementary Movie 5). This embryo was also retrieved after imaging, hatched successfully, and developed into a fertile adult.

Among the four embryos that reached adulthood (embryos 1–3 and 5), marker assessment enabled retrospective genotype assignment (Figure 3, gray rectangles). The first individual lacked both markers and was therefore classified as wild type, consistent with normal development and equivalence to embryos of the reporter subline alone (Strobl et al., 2025). The second and third individuals carried only mCe or mVe, respectively, and were thus identified mono-allelic knockouts for *Zen1*. Their normal development supports the conclusion that *Zen1* is haplosufficient. In contrast, the fifth individual carried both mCe and mVe and was thus identified as a bi-allelic knockout for *Zen1*, consistent with the observed aberrant embryonic phenotype.

Together, these results demonstrate that our four-marker strategy enables reliable organism-level identification of knockout zygosity in a homozygous reporter background, thereby allowing unambiguous correlation of genotype with morphogenetic outcome.

## Discussion

### Summary and conclusion

In this study, we established a modular marker-based genotyping approach that enables organism-level discrimination of knockout zygosity in diploid model organisms and demonstrated its utility for functional analysis during embryogenesis. By introducing spectrally distinct fluorescent transformation markers into insertional mutagenesis approaches, we showed that bi-allelic knockout individuals can be reliably identified and analyzed within mixed cohorts without reliance on workload-intensive molecular methods.

Within the semi-random insertional mutagenesis framework, we identified an adult ‘terminal defect’ phenotype. A remotely comparable condition has been reported following RNAi-mediated knockdown of the *cuticle protein* gene (*CPR27*, NCBI Gene ID: 660347), where structural defects of the cuticle were implicated (Noh et al., 2015). Further, targeted disruption of *Zen1* revealed a semi-lethal loss-of-function condition associated with morphogenetic defects consistent with previously described knockdown phenotypes. Long-term live imaging provided initial mechanistic insight into developmental processes that may underlie reduced viability in respective knockout individuals. Since the present study was designed as a methodological proof-of-principle, detailed functional characterization of both the semi-random insertional phenotype and the *Zen1* loss-of-function condition – including quantitative assessment of penetrance and broader phenotypic analysis – was beyond its scope and will be addressed in future studies. Together, these findings underscore both the developmental relevance of the affected genes and the utility of genotype-resolved live imaging enabled by our marker-based framework.

### Advantages, limitations, and perspective of our approach

The key advantage of this approach is that it preserves the gold-standard framework of gene knockout analysis in diploid organisms while enabling straightforward organism-level genotype discrimination. The four-marker configuration further allows modular combination of independent genetic modifications and thus supports iterative analytical workflows. For example, broadly informative reporters can be used for initial phenotypic characterization, followed by targeted refinement using more specific reporters introduced through repeated application of the crossing scheme. In the case of *Zen1*, a logical next step would be generation of a second knockout-reporter subline using an actin nanobody reporter (Strobl et al., 2025) to directly analyze actomyosin cable dynamics during serosa window closure and dorsal closure.

At the same time, the current implementation has limitations. Although site-specific insertion has been demonstrated in *Tribolium* (Gilles et al., 2015), efficient insertion of larger constructs remains technically challenging. While integrations of ∼1200 bp were achieved, a larger construct (∼2600 bp) containing two markers arranged within interwoven Lox site pairs could not be introduced (data not shown), necessitating two independent rounds of injection. Improvements in HDR efficiency – for example through optimization of delivery strategies or modulation of DNA repair pathways (Chu et al., 2015; Maruyama et al., 2015) – may help overcome this constraint.

A second limitation is that the current marker configuration permits convenient genotype determination only after hatching, denying assessment for embryos that fail to develop further. Future refinements should therefore focus on marker architectures enabling zygosity assessment shortly after nascence and prior to live imaging. Dual-purpose expression cassettes functioning simultaneously as imaging reporters and genotype markers represent a promising strategy. Early-stage genotype identification would enable targeted embryo selection, ensuring balanced sampling across genotypes or prioritization of homozygous knockouts once sufficient controls have been obtained.

Taken together, organism-level genotyping has the potential to develop into a scalable experimental framework for dissecting morphogenetic functions of essential developmental genes by enabling efficient generation, identification and tracking of defined knockout**-**reporter genotypes.

## Methods

### *Tribolium castaneum* strains, rearing, and crossing

In this study, the *Tribolium castaneum* (red flour beetle, NCBI Taxonomy ID: 7070) Plain-White-As-Snow background strain (Strobl et al., 2018) was used, which carries fully unpigmented eyes and thus facilitates visual detection of fluorescent eye-specific transformation markers. Beetles were reared under standard conditions as described previously (Strobl et al., 2025). For single-pair matings, one female-male pair (and five backup pairs whenever possible) were placed in small glass vials containing 1.5 g of growth medium (i.e. full grain wheat flour (SP061036, Demeter) supplemented with 5% (wt/wt) inactive dry yeast (62-106, Flystuff)).

### Germline transformation and subline establishment routine

Germline transformation was, in principle, done as described previously (Lorenzen et al., 2003). Embryos were collected from the parental (P/F0) wild-type culture over a 2-h interval and allowed to rest for 1–2 h at room temperature (23°C ± 1°C). Embryos were then aligned on microscopy slides and injected using a microinjector (FemtoJet, Eppendorf) and 0.7-mm outer-diameter capillaries (Femtotips II, Eppendorf) at an injection pressure of ∼500 hPa. Plasmids were injected in standard buffer (5 mM KCl, 1 mM KH_2_PO_4_ in ddH_2_O, pH 8.8). After hatching, larvae were collected and reared individually in single wells of 24-well plates.

### Semi-random insertion

The piggyBac-based donor plasmid used for semi-random integration was a derivate of the pAVOIAF{#1–#2–#3–#4} plasmid (Strobl et al., 2018) carrying fluorescent eye markers (Berghammer et al., 1999) based on mOrange2 (Shaner et al., 2008) and mVenus (Nagai et al., 2002) in reverse orientation in the #3 and #4 slots. Injections were performed with a mixture containing ∼500 ng/µl of the donor plasmid and ∼400 ng/µl of the pATub’piggyBac helper plasmid (Strobl et al., 2018) encoding a piggyBac transposase under control of the endogenous *tubulin alpha 1-like protein* promoter (Siebert et al., 2008).

### Site-specific integration

The sequence of the *zerknüllt 1* gene (*Zen1*, NCBI Gene ID: 641533) was amplified by polymerase chain reaction (forward primer 5’-CGCGAAAATGGCTTTCTTAAGGGG-3’ and reverse primer 5’-CTCGACGGGTGGTTTTGGCG-3’) from gDNA extracted from adults of the PWAS background strain. The amplicon was sequenced, and a suitable target region for CRISPR/Cas9-mediated induction of a double-strand break was selected based on (i) efficiency predictions generated with CHOPCHOP (Labun et al., 2019) and (ii) the number and likelihood of potential off-target sites identified with Cas-OFFinder (Bae et al., 2014). The homology-directed repair donor plasmids were derived from the commercial pGEM-T Easy plasmid (A1360, Promega) and contained (i) a 950-bp 5’ homology arm identical to the genomic sequence upstream of the double-strand break, (ii) either mVe or mCe, and (iii) a 950 bp 3’ homology arm identical to the genomic sequence upstream of the double-strand break. The gRNA-expressing helper plasmid targeting *zerknüllt 1* was based on the p(U6b-BsaI-gRNA) plasmid (Gilles et al., 2015) and carried 5’-GAGCGGGAATTCCACCAC-3’ as the guide sequence. Injections were performed with a mixture containing ∼300 ng/µl of the homology-directed repair donor plasmid, ∼300 ng/µl of the gRNA-expressing helper plasmid, and ∼300 ng/µl of the pHelper-TC{ATub’NLS-Cas9-NLS) helper plasmid encoding a nuclear-localized Cas9 under control of the endogenous *tubulin alpha 1-like protein* promoter (Siebert et al., 2008).

### Progeny genotype scoring and statistics

Progeny genotypes were scored during the pupal stage. Experimentally obtained marker phenotype distributions were compared to the theoretical Mendelian ratios using a chi-square goodness-of-fit test. Respective *p* values were obtained by Monte Carlo simulation under the corresponding multinomial null model (10^5^ simulations per subline and generation), conditioned on the total number of scored descendants. Python scripts for the statistical calculations can be downloaded at https://github.com/BugCube/Kraemer2026B-Statistics.

### Fluorescence microscopy

Eye markers were imaged using the filter and beam splitter combinations listed in Supplementary Table 4. Light sheet fluorescence microscopy was implemented using a sample chamber-based digital scanned laser light sheet fluorescence microscope (DSLM) (Keller and Stelzer, 2010). A 488 nm / 20 mW diode laser (PhoxX 488-20, Omicron Laserprodukte GmbH) with a 488 nm cleanup filter (xX.F488, Omicron Laserprodukte GmbH) served as the illumination light source. Excitation was performed through a 2.5× NA 0.06 EC Epiplan-Neofluar objective (422320-9900-000, Carl Zeiss AG) while the emission was collected through a 10× NA 0.3 W N-Achroplan objective (420947-9900-000, Carl Zeiss AG). A 525/50 single-band bandpass filter (FF03-525/50-25, Semrock/AHF Analysentechnik AG) and a high-resolution charge-coupled device camera (Clara, Andor) were used for detection. Three micro-translation stages (M-111.2DG, Physik Instrumente GmbH & Co KG) and one precision rotation stage (M-116.DG, Physik Instrumente GmbH & Co KG) were used for sample translation and rotation. The microscope is operated using custom-developed software. Embryo collection, preparation, and mounting were performed essentially as described previously (Ratke et al., 2020). In brief, a transient embryo collection culture was set up using ∼200 (mCe; mC/mC) hemi-heterozygous virgin females and ∼200 (mVe; mC/mC) hemi-heterozygous virgin males. Embryos were collected for 1 h and incubated for 15 h (both at room temperature) to reach the end of blastoderm formation prior to preparation. In the DSLM, axial image (*z*) stacks were acquired in one fluorescence channel along four directions in 90° steps with an interval of 30 min. Embryos were illuminated with a laser power of 135 µW during a 50 ms exposure time window of the camera. All *z* stacks were recorded with a lateral voxel pitch of 0.645 µm and an axial voxel pitch of 2.58 µm (i.e. a pitch ratio of 1:4). Once imaging was completed, embryos were retrieved from the microscope chamber as described previously (Strobl et al., 2015).

### Image processing

Image processing was performed using *Fiji* (Schindelin et al., 2012) (based on *ImageJ* (Schneider et al., 2012), Version 1.53f) and *Mathematica* (Version 13.3.0.0) (Wolfram Research, 2023). Firstly, in *Fiji*, *z* maximum projections were calculated for all *z* stacks and concatenated to time (*t*) stacks. Secondly, using a custom *Mathematica* program (Strobl et al., 2025), *z* and *t* stacks of directions in which embryos were tilted were rotated around *z* to align the anterior-posterior axes of the embryos with the vertical axes of the images (pixels were resampled using a Gaussian kernel) and cropped to remove background. Thirdly, in *Fiji*, the *t* stacks for all directions of a dataset were combined to horizontal t stack montages, which were subjected to histogram matching (Bleach Correction → Histogram Matching) to equalize the signal intensity over the entire time course and then adjusted in brightness and contrast.

## Acknowledgements

The authors thank Ernst H.K. Stelzer for his generous support, including resources and scientific guidance, Sven Plath for technical assistance, and the Goethe University Frankfurt Center for Advanced Light Microscopy (FCAM) core facility for its important contributions to this work.

## Supplementary Figures

**Supplementary Figure 1.**
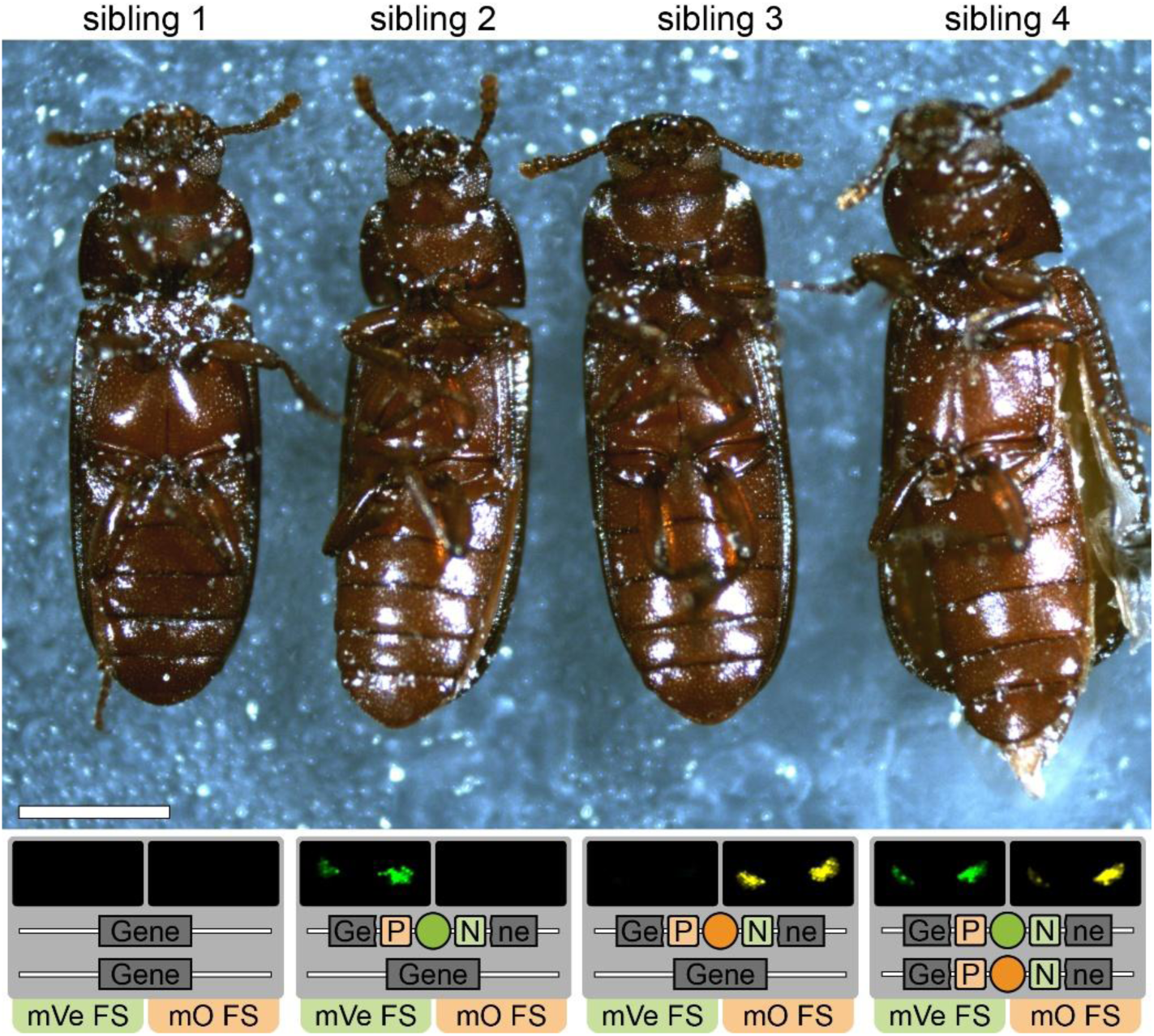
| Marker-based identification of zygosity in females. Four female siblings from the #5 subline derived from a cross between (mVe) and (mO) hemizygotes. Gray boxes show visual markers (pseudo-colored) and inferred genotypes. As for the males, the sibling carrying both mVe and mO, exhibits a pronounced ‘terminal defect’ phenotype. All specimens were alive at the time of imaging. P, LoxP site; N, LoxN site; FS, filter set.

**Supplementary Figure 2.**
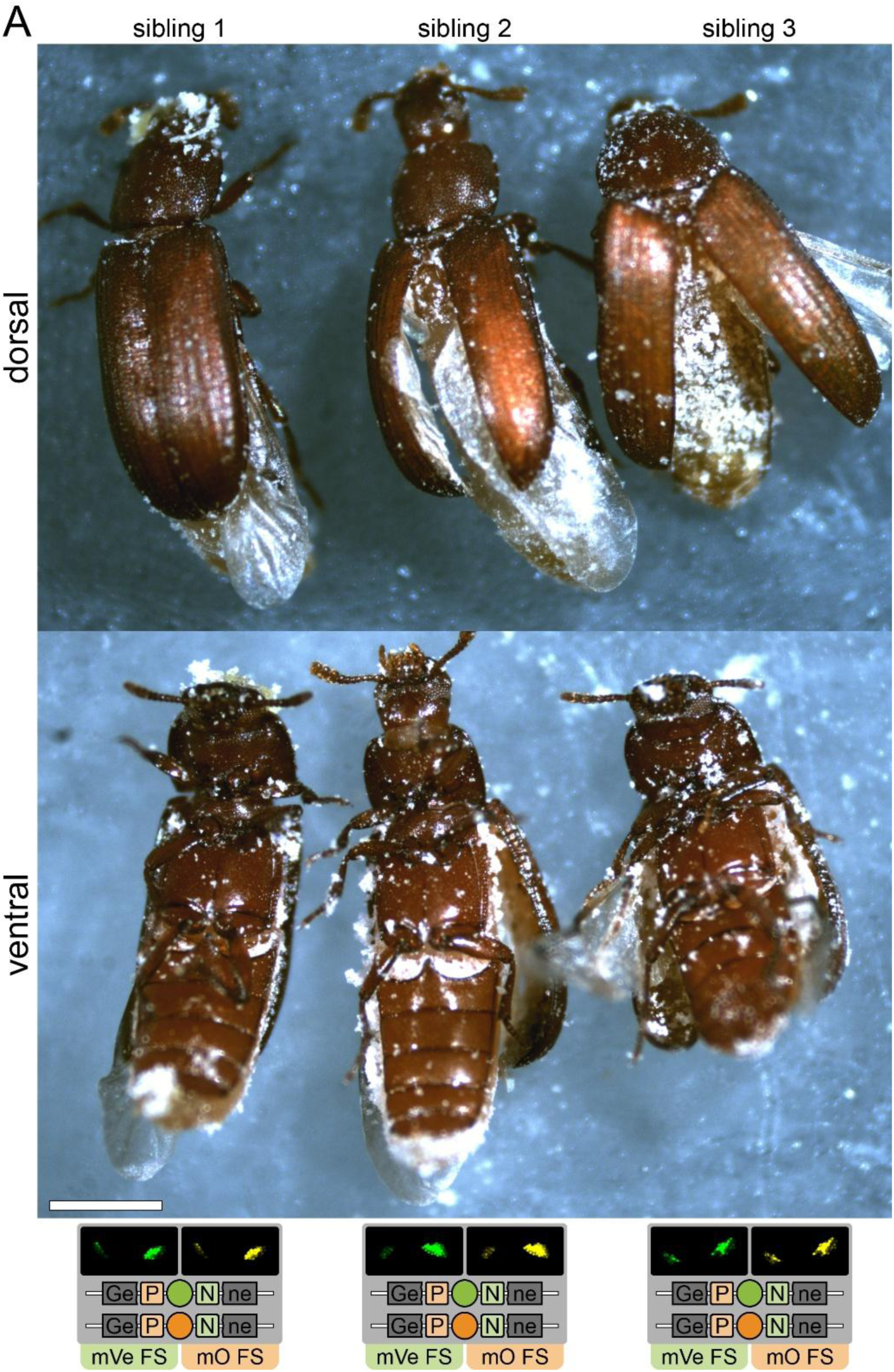
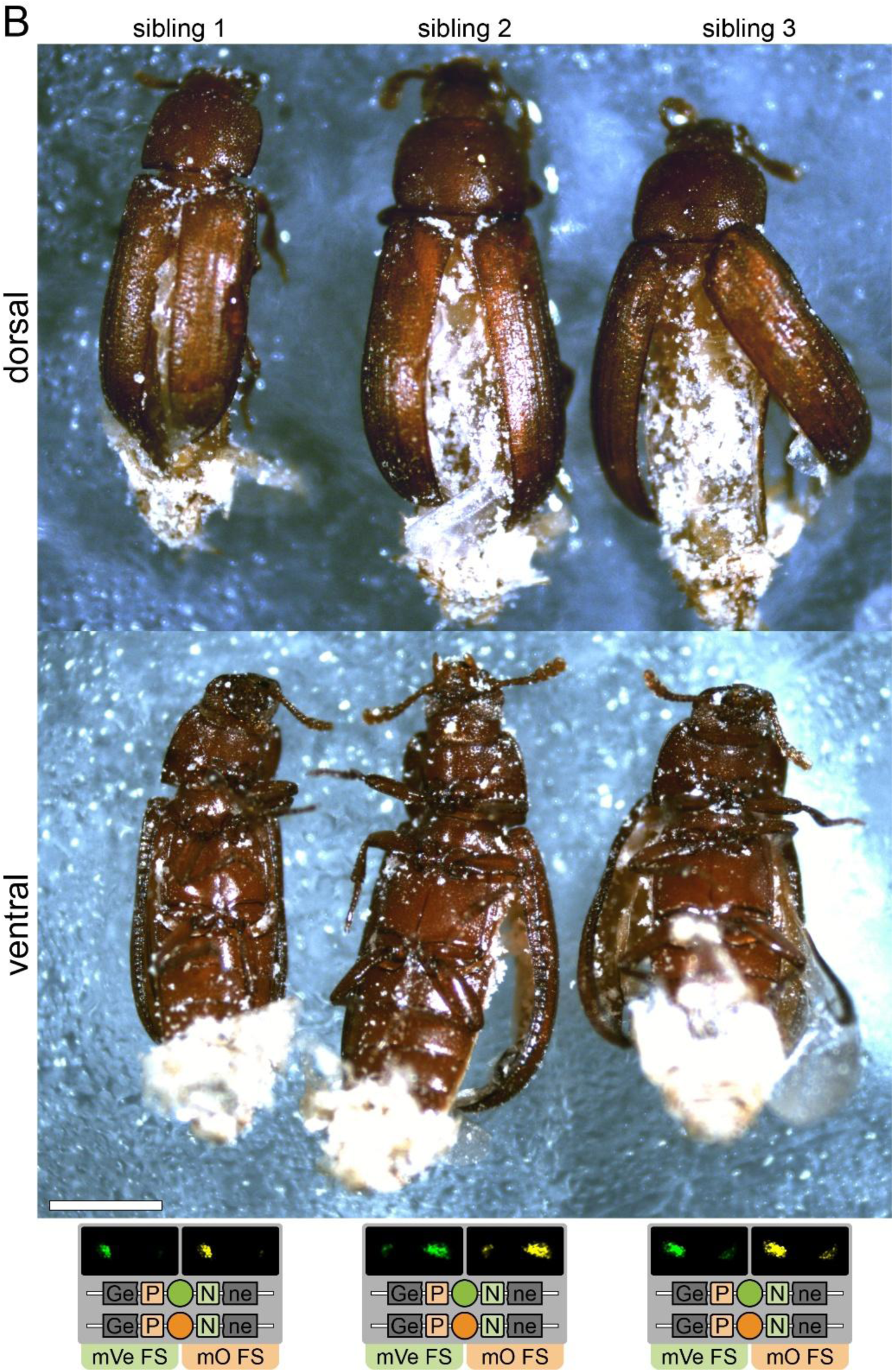
| Phenotypic variance of the ‘terminal defect’ phenotype in the #5 subline. **(a)** Three male siblings carrying both markers shown in dorsal and ventral views. Gray boxes show marker phenotypes (ventral view, pseudo-colored) and inferred genotypes. All individuals were alive at the time of imaging. **(b)** Three female carrying both markers shown in dorsal and ventral views. Gray boxes show visual markers (ventral view). All specimens were alive at the time of imaging. P, LoxP site; N, LoxN site; FS, filter set.

**Supplementary Figure 3.**
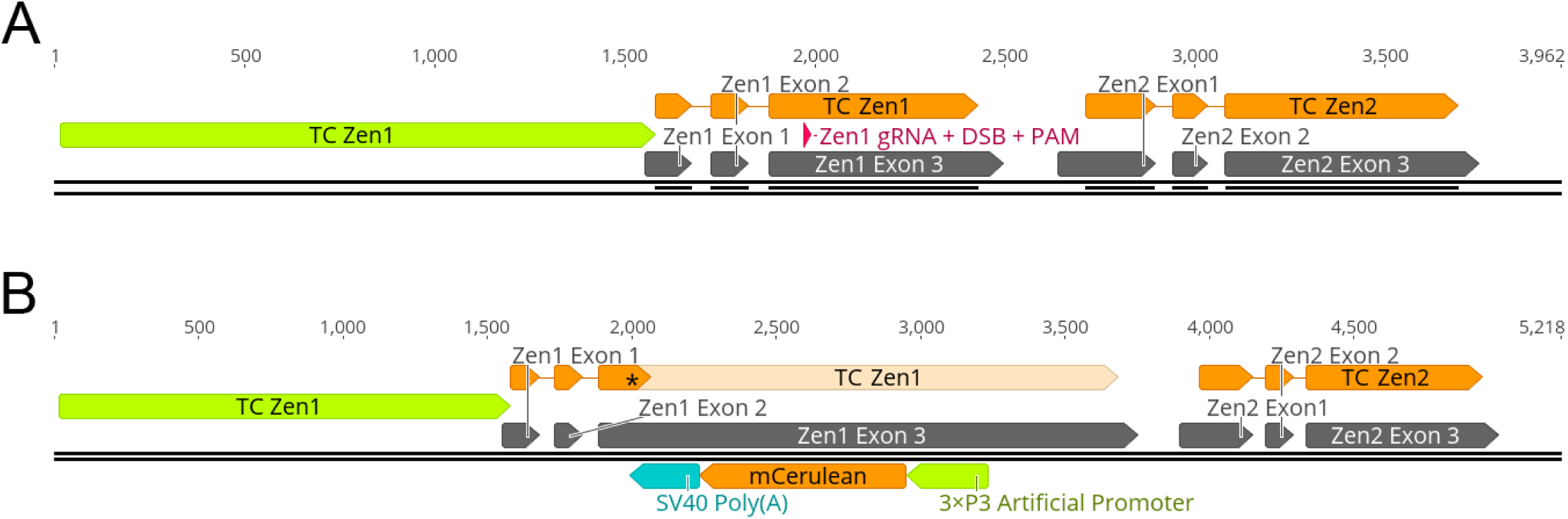
| CRISPR/Cas9-based gene editing approach to insert fluorescent eye markers into the *Zen1* locus. Please note that the scheme illustrates the procedure for mCe; the same procedure was also used for mVe. **(A)** Schematic of the *Zen1* locus in wild-type *Tribolium castaneum* (TC) shown alongside its paralogue *zerknüllt 2* (*Zen2*; NCBI Gene ID: 652932) for reference. The gene comprises three exons (dark gray) that are transcribed into a single mRNA containing one open-reading frame (orange) encoding a 247-amino acid protein. The indicated regulatory/promoter region (green arrow) was previously used to generate a fluorescent reporter line roughly recapitulating *Zen1* expression (Strobl et al., 2018). The red arrowhead indicates the guide RNA target site, pointing to the position of the Cas9-induced double-strand break (DSB) and the protospacer adjacent motif (PAM). **(B)** Schematic of the HDR-mediated integration-based disruption of *Zen1*. The marker, consisting of (i) the eye-specific artificial 3×P3 promoter (Berghammer et al., 1999), (ii) the mCerulean coding sequence, and (iii) the SV40 Poly(A), was inserted in reverse orientation into the third exon. This integration truncates the *Zen1* coding sequence (orange and light orange), which results in a disrupted protein consisting of the first 100 amino acids followed by five missense residues and a premature stop codon.

## Supplementary Tables

**Supplementary Table 1.**
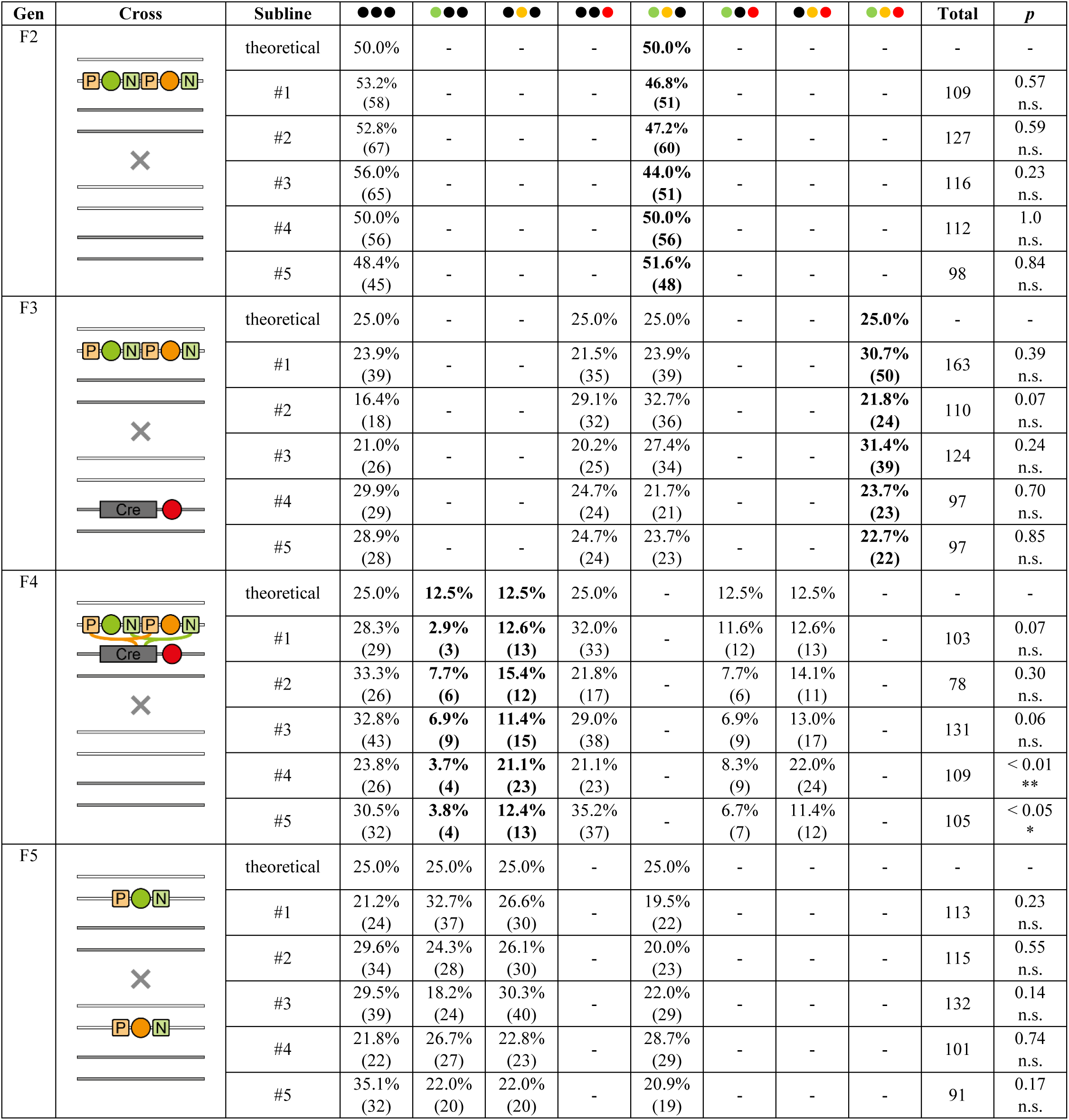
| F2 to F6 crossing results for five transgenic sublines generated via transposon-based semi-random insertional mutagenesis. Progeny genotype frequencies were compared to the theoretical Mendelian ratios (F2, F3, and F5) or a combined expectation incorporation both Mendelian segregation and an unbiased (i.e. 1:1) Lox site pair choice ratio of the Cre recombinase (F4) using a chi-square goodness-of-fit statistic. Percentages and numbers indicate observed genotype frequencies and absolute progeny scores, respectively. No significant deviations were detected in the F2, F3, and F5. Significant deviations observed in the F4 for two of the sublines (#4 and #5) are likely attributable to biased Lox site pair choice, consistent with previous observations (Strobl et al., 2023). P, LoxP site; N, LoxN site; Cre, expression cassette for the Cre recombinase; n.s., not significant.

| Gen | Cross | Subline | ●●● | ●●● | ●●● | ●●● | ●●● | ●●● | ●●● | ●●● | Total | p |
| --- | --- | --- | --- | --- | --- | --- | --- | --- | --- | --- | --- | --- |
| F2 |  | theoretical | 50.0% | - | - | - | 50.0% | - | - | - | - | - |
|  |  | #1 | 53.2%<br>(58) | - | - | - | 46.8%<br>(51) | - | - | - | 109 | 0.57<br>n.s. |
|  |  | #2 | 52.8%<br>(67) | - | - | - | 47.2%<br>(60) | - | - | - | 127 | 0.59<br>n.s. |
|  |  | #3 | 56.0%<br>(65) | - | - | - | 44.0%<br>(51) | - | - | - | 116 | 0.23<br>n.s. |
|  |  | #4 | 50.0%<br>(56) | - | - | - | 50.0%<br>(56) | - | - | - | 112 | 1.0<br>n.s. |
|  |  | #5 | 48.4%<br>(45) | - | - | - | 51.6%<br>(48) | - | - | - | 98 | 0.84<br>n.s. |
| F3 |  | theoretical | 25.0% | - | - | 25.0% | 25.0% | - | - | 25.0% | - | - |
|  |  | #1 | 23.9%<br>(39) | - | - | 21.5%<br>(35) | 23.9%<br>(39) | - | - | 30.7%<br>(50) | 163 | 0.39<br>n.s. |
|  |  | #2 | 16.4%<br>(18) | - | - | 29.1%<br>(32) | 32.7%<br>(36) | - | - | 21.8%<br>(24) | 110 | 0.07<br>n.s. |
|  |  | #3 | 21.0%<br>(26) | - | - | 20.2%<br>(25) | 27.4%<br>(34) | - | - | 31.4%<br>(39) | 124 | 0.24<br>n.s. |
|  |  | #4 | 29.9%<br>(29) | - | - | 24.7%<br>(24) | 21.7%<br>(21) | - | - | 23.7%<br>(23) | 97 | 0.70<br>n.s. |
|  |  | #5 | 28.9%<br>(28) | - | - | 24.7%<br>(24) | 23.7%<br>(23) | - | - | 22.7%<br>(22) | 97 | 0.85<br>n.s. |
| F4 |  | theoretical | 25.0% | 12.5% | 12.5% | 25.0% | - | 12.5% | 12.5% | - | - | - |
|  |  | #1 | 28.3%<br>(29) | 2.9%<br>(3) | 12.6%<br>(13) | 32.0%<br>(33) | - | 11.6%<br>(12) | 12.6%<br>(13) | - | 103 | 0.07<br>n.s. |
|  |  | #2 | 33.3%<br>(26) | 7.7%<br>(6) | 15.4%<br>(12) | 21.8%<br>(17) | - | 7.7%<br>(6) | 14.1%<br>(11) | - | 78 | 0.30<br>n.s. |
|  |  | #3 | 32.8%<br>(43) | 6.9%<br>(9) | 11.4%<br>(15) | 29.0%<br>(38) | - | 6.9%<br>(9) | 13.0%<br>(17) | - | 131 | 0.06<br>n.s. |
|  |  | #4 | 23.8%<br>(26) | 3.7%<br>(4) | 21.1%<br>(23) | 21.1%<br>(23) | - | 8.3%<br>(9) | 22.0%<br>(24) | - | 109 | < 0.01<br>** |
|  |  | #5 | 30.5%<br>(32) | 3.8%<br>(4) | 12.4%<br>(13) | 35.2%<br>(37) | - | 6.7%<br>(7) | 11.4%<br>(12) | - | 105 | < 0.05<br>* |
| F5 |  | theoretical | 25.0% | 25.0% | 25.0% | - | 25.0% | - | - | - | - | - |
|  |  | #1 | 21.2%<br>(24) | 32.7%<br>(37) | 26.6%<br>(30) | - | 19.5%<br>(22) | - | - | - | 113 | 0.23<br>n.s. |
|  |  | #2 | 29.6%<br>(34) | 24.3%<br>(28) | 26.1%<br>(30) | - | 20.0%<br>(23) | - | - | - | 115 | 0.55<br>n.s. |
|  |  | #3 | 29.5%<br>(39) | 18.2%<br>(24) | 30.3%<br>(40) | - | 22.0%<br>(29) | - | - | - | 132 | 0.14<br>n.s. |
|  |  | #4 | 21.8%<br>(22) | 26.7%<br>(27) | 22.8%<br>(23) | - | 28.7%<br>(29) | - | - | - | 101 | 0.74<br>n.s. |
|  |  | #5 | 35.1%<br>(32) | 22.0%<br>(20) | 22.0%<br>(20) | - | 20.9%<br>(19) | - | - | - | 91 | 0.17<br>n.s. |

**Supplementary Table 2.** | F2 to F6 crossing results for all three replicates deriving from the F2 founder carrying mCe. Progeny genotype frequencies were compared to the theoretical Mendelian ratios. Percentages and numbers indicate observed genotype frequencies and absolute progeny scores, respectively. No significant deviations were detected. H2°NB-mE, expression cassette for mEmerald-labeled anti-histone nanobodies.

| Gen | Cross | Replicate | ●●● | ●●● | ●●● | ●●● | ●●● | ●●● | ●●● | ●●● | Total | *p* |
| --- | --- | --- | --- | --- | --- | --- | --- | --- | --- | --- | --- | --- |
| F3 |  | theoretical | 50.0% | **50.0%** | - | - | - | - | - | - | - | - |
|  |  | 1 | 47.4% (54) | **52.6% (60)** | - | - | - | - | - | - | 114 | 0.64 (n.s.) |
|  |  | 2 | 51.3% (39) | **48.7% (37)** | - | - | - | - | - | - | 76 | 0.91 (n.s.) |
|  |  | 3 | 44.4% (16) | **55.6% (20)** | - | - | - | - | - | - | 36 | 0.62 (n.s.) |
| F4 |  | theoretical | - | - | 50.0% | - | **50.0%** | - | - | - | - | - |
|  |  | 1 | - | - | 52.2% (72) | - | **47.8% (66)** | - | - | - | 138 | 0.67 (n.s.) |
|  |  | 2 | - | - | 50.0% (45) | - | **50.0% (45)** | - | - | - | 90 | 1.00 (n.s.) |
|  |  | 3 | - | - | 53.3% (48) | - | **46.7% (42)** | - | - | - | 90 | 0.60 (n.s.) |
| F5 |  | theoretical | - | - | - | 25.0% | - | 25.0% | 25.0% | **25.0%** | - | - |
|  |  | 1 | - | - | - | 30.0% (12) | - | 22.5% (9) | 25.0% (10) | **22.5% (9)** | - | 0.94 (n.s.) |
|  |  | 2 | - | - | - | 25.6% (20) | - | 24.4% (19) | 28.2% (22) | **21.8% (17)** | - | 0.89 (n.s.) |
|  |  | 3 | - | - | - | 23.2% (17) | - | 27.4% (20) | 24.7% (18) | **24.7% (18)** | - | 0.97 (n.s.) |
| F6 |  | theoretical | - | - | 12.5% | 12.5% | 12.5% | **12.5%** | 25.0% | 25.0% | - | - |
|  |  | 1 | - | - | 12.4% (15) | 10.7% (13) | 11.6% (14) | **14.9% (18)** | 25.6% (31) | 24.8% (30) | 121 | 0.95 (n.s.) |
|  |  | 2 | - | - | 13.9% (10) | 12.5% (9) | 8.3% (6) | **12.5% (9)** | 25.0% (18) | 27.8% (20) | 72 | 0.97 (n.s.) |
|  |  | 3 | - | - | 9.7% (9) | 17.2% (16) | 14.0% (13) | **10.7% (10)** | 28.0% (26) | 20.4% (19) | 93 | 0.93 (n.s.) |
| F7C |  | theoretical | - | - | - | 50.0% | - | 50.0% | - | - | - | - |
|  |  | 1 | - | - | - | 47.6% (30) | - | 53.4% (33) | - | - | 63 | 0.80 (n.s.) |
|  |  | 2 | - | - | - | 51.3% (40) | - | 48.7% (38) | - | - | 78 | 0.91 (n.s.) |
|  |  | 3 | - | - | - | 47.3% (53) | - | 52.7% (59) | - | - | 112 | 0.64 (n.s.) |

**Supplementary Table 3.** | F2 to F6 crossing results for all three replicates deriving from the F2 founder carrying mVe. Progeny genotype frequencies were compared to the theoretical Mendelian ratios. Percentages and numbers indicate observed genotype frequencies and absolute progeny scores, respectively. No significant deviations were detected. H2°NB-mE, expression cassette for mEmerald-labeled anti-histone nanobodies.

| Gen | Cross | Replicate | ●●● | ●●● | ●●● | ●●● | ●●● | ●●● | ●●● | ●●● | Total | p |
| --- | --- | --- | --- | --- | --- | --- | --- | --- | --- | --- | --- | --- |
| F3 |  | theoretical | 50.0% | **50.0%** | - | - | - | - | - | - | - | - |
|  |  | 1 | 46.7%<br>(42) | **53.3%**<br>**(48)** | - | - | - | - | - | - | 90 | 0.60<br>(n.s.) |
|  |  | 2 | 54.1%<br>(46) | **45.1%**<br>**(39)** | - | - | - | - | - | - | 85 | 0.51<br>(n.s.) |
|  |  | 3 | 48.5%<br>(50) | **51.5%**<br>**(53)** | - | - | - | - | - | - | 103 | 0.84<br>(n.s.) |
| F4 |  | theoretical | - | - | 50.0% | - | **50.0%** | - | - | - | - | - |
|  |  | 1 | - | - | 51.6%<br>(66) | - | **48.4%**<br>**(62)** | - | - | - | 128 | 0.79<br>(n.s.) |
|  |  | 2 | - | - | 48.2%<br>(40) | - | **51.8%**<br>**(43)** | - | - | - | 83 | 0.83<br>(n.s.) |
|  |  | 3 | - | - | 52.1%<br>(61) | - | **47.9%**<br>**(56)** | - | - | - | 117 | 0.71<br>(n.s.) |
| F5 |  | theoretical | - | - | - | 25.0% | - | 25.0% | 25.0% | **25.0%** | - | - |
|  |  | 1 | - | - | - | 19.7%<br>(14) | - | 25.4%<br>(18) | 29.5%<br>(21) | **25.4%**<br>**(18)** | 71 | 0.71<br>(n.s.) |
|  |  | 2 | - | - | - | 24.1%<br>(26) | - | 28.7%<br>(31) | 22.2%<br>(24) | **25.0%**<br>**(27)** | 108 | 0.83<br>(n.s.) |
|  |  | 3 | - | - | - | 23.8%<br>(20) | - | 26.2%<br>(22) | 23.8%<br>(20) | **26.2%**<br>**(22)** | 84 | 0.98<br>(n.s.) |
| F6 |  | theoretical | - | - | 12.5% | 12.5% | 12.5% | **12.5%** | 25.0% | 25.0% | - | - |
|  |  | 1 | - | - | 11.6%<br>(10) | 14.0%<br>(12) | 10.5%<br>(9) | **14.0%**<br>**(12)** | 23.2%<br>(20) | 26.7%<br>(23) | 86 | 0.98<br>(n.s.) |
|  |  | 2 | - | - | 12.7%<br>(14) | 12.7%<br>(14) | 15.5%<br>(17) | **12.7%**<br>**(14)** | 21.8%<br>(24) | 24.6%<br>(27) | 110 | 0.95<br>(n.s.) |
|  |  | 3 | - | - | 9.8%<br>(9) | 15.2%<br>(14) | 13.0%<br>(12) | **14.1%**<br>**(13)** | 27.2%<br>(25) | 20.7%<br>(19) | 92 | 0.84<br>(n.s.) |
| F7C |  | theoretical | - | - | - | 50.0% | - | 50.0% | - | - | - | - |
|  |  | 1 | - | - | - | 53.7%<br>(44) | - | 46.3%<br>(38) | - | - | 82 | 0.58<br>(n.s.) |
|  |  | 2 | - | - | - | 45.5%<br>(51) | - | 54.5%<br>(61) | - | - | 112 | 0.39<br>(n.s.) |
|  |  | 3 | - | - | - | 50.5%<br>(48) | - | 49.5%<br>(47) | - | - | 95 | 1.00<br>(n.s.) |

**Supplementary Table 4.**
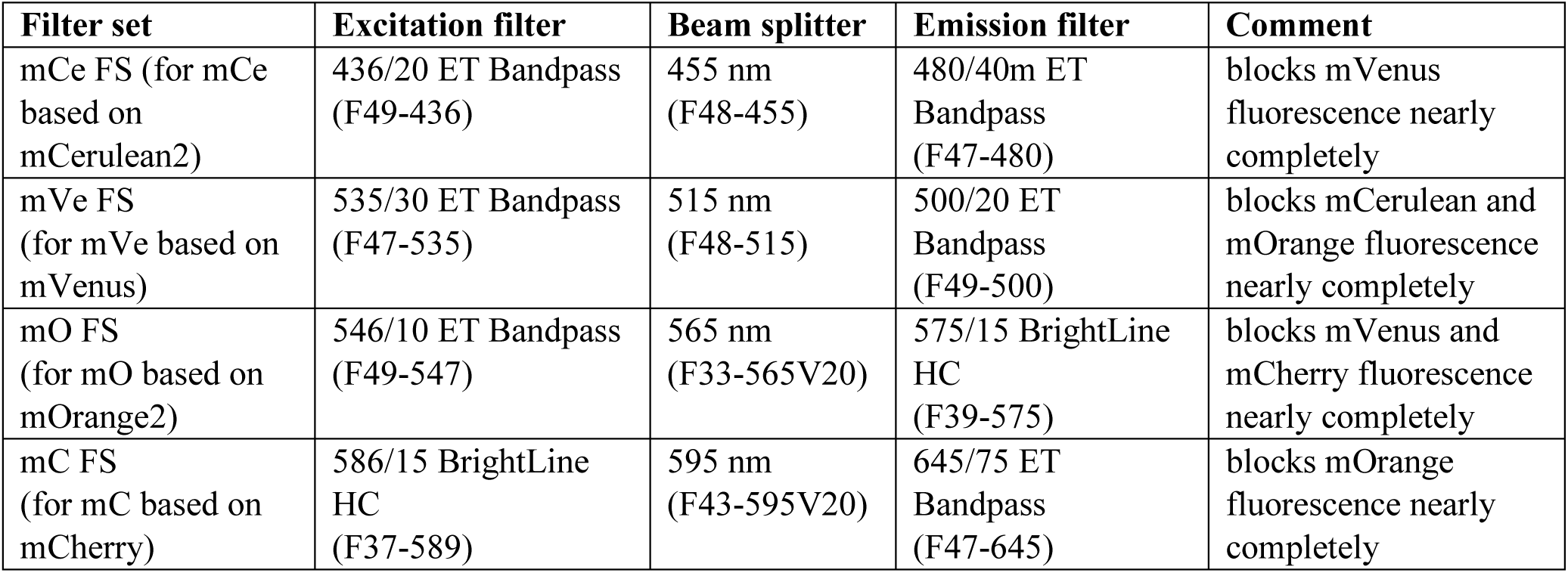
| Fluorescence filter sets for fluorescence eye marker documentation. All components were obtained from AHF Analysentechnik, Germany. FS, filter set.

## Supplementary Movies

**Supplementary Movie 1 | Long-term live imaging of a *Tribolium castaneum* embryo that showed normal development and hatched after retrieval.** The embryo originated from the F7 (mCe; mC/mC) × (mVe; mC/mC) cross and was thus homozygous for transgene expressing mEmerald-labeled nanobodies against histones H2A and H2B. The specimen is shown along four orientations spaced 90° apart and was imaged over 122:30 h with a 30-min interval. ZH, *z* maximum projection with intensity adjustments and histogram correction.

**Supplementary Movie 2 | Long-term live imaging of a *Tribolium castaneum* embryo that showed normal development and hatched after retrieval.** The embryo originated from the F7 (mCe; mC/mC) × (mVe; mC/mC) cross and was thus homozygous for transgene expressing mEmerald-labeled nanobodies against histones H2A and H2B. The specimen is shown along four orientations spaced 90° apart and was imaged over 101:00 h with a 30-min interval. ZH, *z* maximum projection with intensity adjustments and histogram correction.

**Supplementary Movie 3 | Long-term live imaging of a *Tribolium castaneum* embryo that showed normal development and hatched after retrieval.** The embryo originated from the F7 (mCe; mC/mC) × (mVe; mC/mC) cross and was thus homozygous for transgene expressing mEmerald-labeled nanobodies against histones H2A and H2B. The specimen is shown along four orientations spaced 90° apart and was imaged over 117:00 h with a 30-min interval. ZH, *z* maximum projection with intensity adjustments and histogram correction.

**Supplementary Movie 4 | Long-term live imaging of a *Tribolium castaneum* embryo that showed aberrational development and did not hatch after retrieval.** The embryo originated from the F7 (mCe; mC/mC) × (mVe; mC/mC) cross and was thus homozygous for transgene expressing mEmerald-labeled nanobodies against histones H2A and H2B. The specimen is shown along four orientations spaced 90° apart and was imaged over 120:00 h with a 30-min interval. ZH, *z* maximum projection with intensity adjustments and histogram correction.

**Supplementary Movie 5 | Long-term live imaging of a *Tribolium castaneum* embryo that showed aberrational development and hatched after retrieval.** The embryo originated from the F7 (mCe; mC/mC) × (mVe; mC/mC) cross and was thus homozygous for transgene expressing mEmerald-labeled nanobodies against histones H2A and H2B. The specimen is shown along four orientations spaced 90° apart and was imaged over 122:30 h with a 30-min interval. ZH, *z* maximum projection with intensity adjustments and histogram correction.

